# Multimodal profiling of chordoma immunity reveals distinct immune contextures

**DOI:** 10.1101/2023.09.14.557579

**Authors:** S. van Oost, D.M. Meijer, M.E. IJsselsteijn, J. Roelands, B. van den Akker, R. van der Breggen, I.H. Briaire-de Bruijn, M. van der Ploeg, P.M. Wijers-Koster, S.B. Polak, W.C. Peul, R.J.P. van der Wal, N.F.C.C. de Miranda, J.V.M.G. Bovee

## Abstract

Chordomas are cancers from the axial skeleton presenting immunological hallmarks of unknown significance. In recent years, some clinical trials demonstrated that chordomas can respond to immunotherapy. We present a comprehensive characterisation of immunological features of 76 chordomas through application of a multimodal approach comprising transcriptional profiling, multidimensional immunophenotyping and TCR profiling. Chordomas generally presented an immune “hot” microenvironment in comparison to other sarcomas, as indicated by the immunologic constant of rejection transcriptional signature. We identified two distinct groups of chordomas based on T cell infiltration. The highly infiltrated group was further characterised by high dendritic cell infiltration and the presence of multicellular immune aggregates in tumours, whereas low T cell infiltration was associated with lower overall cell densities of immune and stromal cells. Interestingly, patients with higher T cell infiltration displayed a more pronounced clonal enrichment of the T cell receptor repertoire compared to those with low T cell counts. Furthermore, we observed that the majority of chordomas maintained HLA class I expression. Our findings shed light on the natural immunity against chordomas. Understanding their immune landscape could guide the development and application of immunotherapies in a tailored manner, ultimately leading to an improved clinical outcome for chordoma patients.

## Introduction

Chordomas are cancers displaying notochordal differentiation that arise predominantly in the axial skeleton at the skull base, the craniocervical and mobile spine, the sacrum, and, occasionally, at extra-axial areas^1^. General treatment for chordomas consists of surgical resection and/or (neo-)adjuvant proton beam radiation. Since most chordomas are indolent tumours, chemotherapy is considered ineffective for the treatment of this disease^2^. The clinical management of chordoma patients is complicated by the anatomical location of the tumours, their often late detection, as well as treatment-related morbidities^2,3^. Aiming at a gross total or en bloc resection involves sacrifice of neurovascular structures with permanent neurological disabilities. As such, disease recurrence is common after treatment and 30-40% of chordoma patients develop metastases^4^. The overall 5-year survival following diagnosis is around 67% and there are currently no curative therapies for chordoma patients with advanced disease^1,5^. This reinforces the need for innovative treatments as local recurrences and distant metastases persist.

Recent findings have provided evidence for the presence of conspicuous immune cell infiltration in chordomas^6,7^ and, encouragingly, clinical responses of chordoma patients to checkpoint blockade immunotherapy targeting CTLA-4 and/or the PD-1/PD-L1 axis have been reported^8–12^. This is an intriguing observation as chordomas display a low mutational burden and are, instead, affected by gross chromosomal alterations^13–15^. Furthermore, PD-L1 expression is rarely encountered in sarcomas, including chordomas^16–18^. Instead, the presence of pre-existing anti-tumour immune responses perceived by the infiltration of cytotoxic T cells in the tumour tissue or by the presence of tertiary lymphoid structures appear to inform on the sensitivity of soft tissue sarcomas to immune checkpoint blockade (ICB)^19,20^. So far, whether tertiary lymphoid structures are predictive of ICB responsiveness has not been investigated in chordomas.

An in-depth immunological characterisation of chordomas and their association with clinical parameters is currently lacking. Here, we applied transcriptomic profiling and multidimensional immunophenotyping by imaging mass cytometry and multispectral immunofluorescence to resolve the immune landscape of chordomas. Chordomas presented hallmarks of anti-tumour immunity and were considered immune “hot” in comparison with other sarcomas. Within our cohort, we identified groups of chordomas with distinct immune contextures, independent from any clinical parameter. Our findings support the rational utilisation of immunotherapeutic approaches, tailored to target these distinct microenvironments in chordoma.

## Results

### Patient characteristics

In this study, two cohorts were established to investigate the immune microenvironment of chordomas. The first cohort included primary tumours of 32 patients and was explored using RNA sequencing (*n* = 20) and imaging mass cytometry (*n* = 32). The median age was 59 years (range 1 - 80 years) and the median follow-up time was 66 months (range 0 - 418 months). In total, the primary tumours in the cohort comprised eight skull base chordomas (25%), seven spinal chordomas (22%) and 17 sacrococcygeal chordomas (53%). The second cohort included primary tumours of an additional 53 patients. The median age of this cohort was 60 years (range 22 – 80) and the median follow-up time was 41 months (range 0 - 182). In total, this cohort comprised 17 skull base (32%), 15 spinal chordomas (28%), 20 sacrococcygeal chordomas (38%) and one extra-axial chordoma of the ribs (2%). The second cohort also comprised patients solely treated with radiotherapy, for which the treatment-naïve biopsies were used. All profiled primary tumour samples were annotated for their anatomical location as well as whether they underwent neoadjuvant radiotherapy. Further patient characteristics of both cohorts are listed in **Table 1**.

**Table 1.** Patient characteristics of chordoma cohorts. This table contains an overview of each cohort separately as well as both cohorts combined.

| Variable | First |  | Second |  | Combined |  |
| --- | --- | --- | --- | --- | --- | --- |
| Total patients, *n* | 32 |  | 53 |  | 85 |  |
| Gender, *n* (%) |  |  |  |  |  |  |
| Female | 15 | (46.9) | 21 | (39.6) | 36 | (42.4) |
| Male | 17 | (53.1) | 32 | (60.4) | 49 | (57.6) |
| Age (years), *median* (range) | 59 | (1-80) | 60 | (22-80) | 60 | (1-80) |
| Follow-up (months), *median* (range) | 66 | (0-418) | 41 | (0-182) | 50 | (0-418) |
| Anatomical location, *n* (%)<sup>a</sup> |  |  |  |  |  |  |
| Skull base (Clivus) | 8 | (25.0) | 17 | (32.1) | 25 | (29.4) |
| Mobile spine (Craniocervical, Cervical, Thoracic, Lumbar) | 7 (0, 1, 4, 2) | (21.9) | 15 (3, 10, 1, 1) | (28.3) | 22 | (25.9) |
| Sacrococcygeal | 17 | (53.1) | 20 | (37.7) | 27 | (31.8) |
| Extra-axial (Costa) | - | - | 1 | (1.9) | 1 | (1.2) |
| Max tumour size (cm), *median* (IQR)<sup>b</sup> | 7 | (4.5) | 6 | (6.1) | 6.1 | (5.6) |
| Skull base (Clivus) | 2.8 | (0.88) | 4.3 | (1.5) | 4 | (2.5) |
| Mobile spine (Craniocervical, Cervical, Thoracic, Lumbar) | 5.5 | (3.4) | 4.9 | (1.4) | 5 | (1.7) |
| Sacrococcygeal | 8 | (4) | 10.5 | (3.6) | 9 | (3.9) |
| Extra-axial (Costa) | - | - | 10 | - | 10 | - |
| Treatment, *n* (%) |  |  |  |  |  |  |
| Radiotherapy alone | - | - | 11 | (20.8) | 11 | (12.9) |
| Surgery alone | 21 | (65.6) | 19 | (35.8) | 40 | (47.1) |
| Surgery & radiotherapy | 11 | (34.4) | 23 | (43.4) | 34 | (40) |
| Other radiotherapy regimens, *n* (%) |  |  |  |  |  |  |
| Monotherapy, conventional | - | - | 1 | (1.9) | 1 | (1.2) |
| Monotherapy, proton beam | - | - | 9 | (17) | 9 | (10.6) |
| Monotherapy, carbon ion | - | - | 1 | (1.9) | 1 | (1.2) |
| Sandwich, proton beam | - | - | 2 | (3.8) | 2 | (2.4) |
| Pre-operative radiotherapy, *n* (%)<sup>c</sup> |  |  |  |  |  |  |
| Conventional | 4 | (12.5) | 2 | (3.8) | 6 | (7.1) |
| Proton beam | 2 | (6.3) | 1 | (1.9) | 3 | (3.5) |
| Post-operative radiotherapy, *n* (%)<sup>c</sup> |  |  |  |  |  |  |
| Conventional | 5 | (15.6) | 7 | (13.2) | 12 | (14.1) |
| Proton beam | - | - | 11 | (20.8) | 11 | (12.9) |
| Patients with recurrent disease, *n* (%)<sup>d</sup> |  |  |  |  |  |  |
| Local recurrence | 16 | (50) | 29 | (54.7) | 45 | (52.9) |
| Metastasis | 6 | (18.8) | 14 | (26.4) | 20 | (23.5) |

<sup>a</sup>anatomical location of the primary tumours; <sup>b</sup>size was not available for 8 and 11 primary tumours in the initial and the additional cohort respectively (4 skull base and 2 mobile spine; 2 skull base, 2 mobile spine and 7 sacrococcygeal);
<sup>c</sup>treatments regarding primary tumours; <sup>d</sup>Patients who experienced recurrent disease during follow-up. With the exception of 3 patients in the additional cohort, metastatic disease followed local recurrence.

### Chordomas present transcriptional hallmarks of anti-cancer immunity

The transcriptomic profiles of the 20 chordomas were compared to those of osteosarcomas (*n* = 7), chondrosarcomas (*n* = 9), undifferentiated soft tissue sarcomas (USTS) (*n* = 13), and myxofibrosarcomas (*n* = 10). All chordoma samples clustered separately from the remaining sarcomas following unsupervised clustering based on the 200 genes with most variable expression among all samples (**Fig. 1A**). Sample clustering was largely driven by features related to the line of differentiation of the different sarcoma samples, with known chordoma-related genes such as *TBXT*, *CD24*, and several keratins enriched in the chordomas^21–23^ (**Fig. 1A**, **Supplementary Table 1**). Moreover, genes associated with the extracellular matrix and adhesion related genes were enriched in chordomas and chondrosarcomas as compared to the other sarcomas (**Fig. 1A**, **Supplementary Table 1**). Interestingly, several immune-related genes, including Human Leukocyte Antigen (HLA) class I, HLA class II, *PLA2G2D* and immunoglobulin genes, were among the most variably expressed genes and were enriched in specific chordomas. Furthermore, differential gene expression analysis revealed enrichment for other immune-related genes in chordomas in comparison with the other sarcomas (**Supplementary Fig. 1A-B**, **Supplementary Table 2**).

**Fig. 1.**
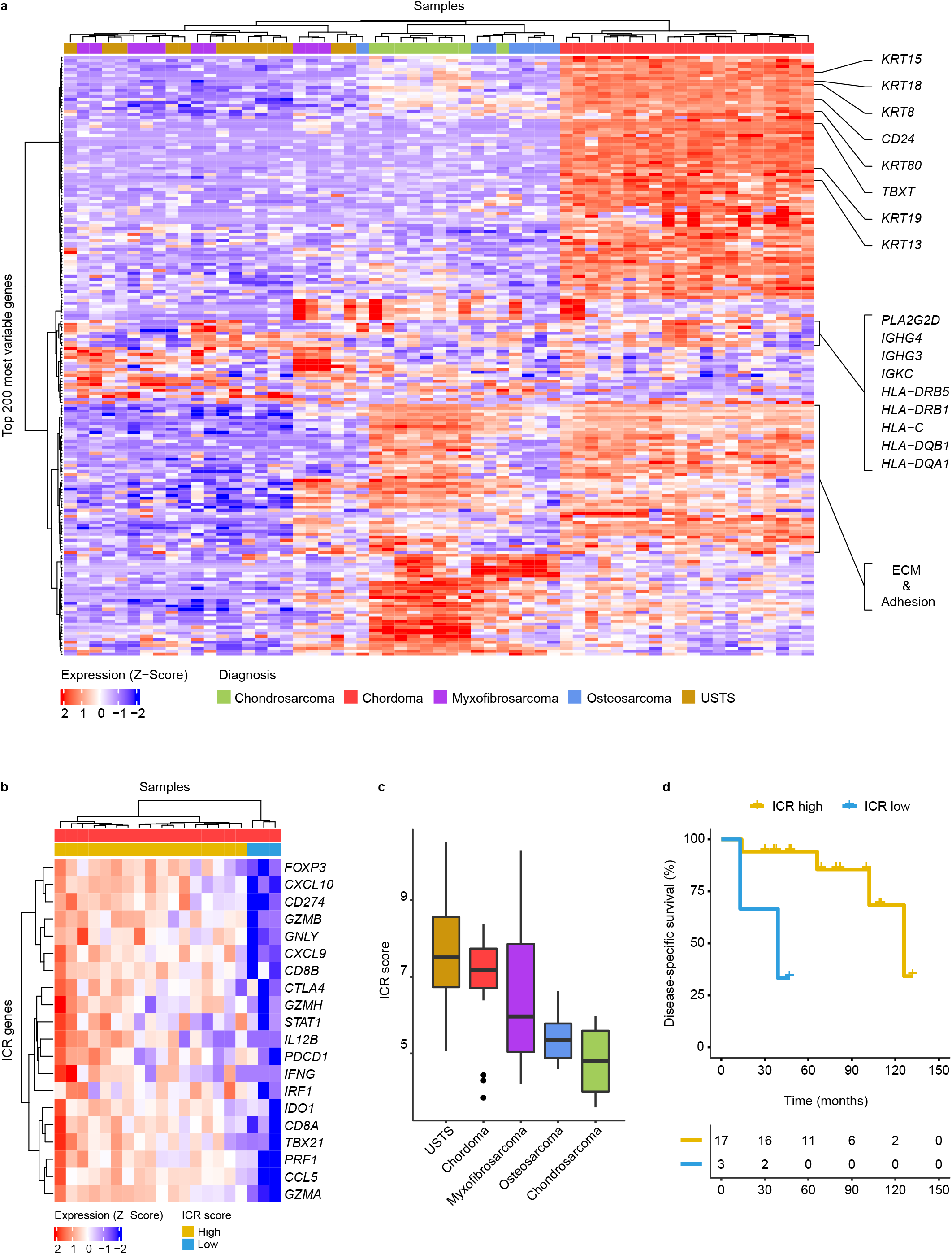
Chordomas display hallmarks of anti-tumour immunity. **a**, Heatmap demonstrating the log2 normalised gene expression Z-Score of the top 200 most variably expressed genes in all sarcoma samples (*n* = 59). Samples are annotated according to their histotype. Genes associated with chordoma samples are highlighted on the right. These genes include chordoma-specific genes such as *TBXT, CD24* and several keratins. In addition, immune-related genes are highlighted, including Human Leukocyte Antigen (*HLA*\*) and immunoglobulin (*IG*\*) genes. **b**, Heatmap presenting the Z-Score for the log_2_ normalised expression of the 20 ICR signature genes in 20 chordoma samples. Based on the expression Z-Score, the samples are classified into two groups through unsupervised hierarchical clustering: ICR high (*n* = 17) and ICR low (*n* = 3). **c**, Boxplots presenting the Immunologic Constant of Rejection (ICR) score per sarcoma subtype in descending order. **d**, Kaplan-Meier curves displaying disease-specific survival for the ICR high and ICR low chordomas. No statistical analysis was performed due to the small sample size. ECM = extracellular matrix; USTS = undifferentiated soft tissue sarcoma.

To substantiate that immune-related processes are enriched in chordomas, we evaluated the expression of genes included in the Immunologic Constant of Rejection (ICR) gene signature to compare between the distinct sarcoma subtypes^24^. This 20 gene signature reflects the activation of type 1 T (Th1) cell signalling, expression of CXCR3/CCR5 chemokine ligands, cytotoxicity and counter-activation of immunoregulatory mechanisms and was previously shown to be an independent prognostic factor for metastasis-free survival in soft tissue sarcomas^25^. Most chordomas are characterised by a high expression of ICR genes (ICR high) (**Fig. 1B**), particularly when comparing the ICR score (average of the 20 ICR genes) with samples from other sarcoma types (**Fig. 1C**). Furthermore, the ICR high immune subtype was associated with improved disease-specific survival in this cohort, albeit only three cases were classified as ICR low (**Fig. 1D**).

### T cell infiltration in chordomas correlates with ICR score and separates patients into distinct groups

To confirm and expand our initial observations, we performed imaging mass cytometry to characterise the immune cell contexture of 32 primary tumours. Twenty-nine cell populations were identified including five T cell phenotypes comprising CD4^+^ T cells, CD8^+^ T cells, Ki-67^+^ T cells, regulatory T cells and (unspecified) T cells including low numbers of CD57^+^ and gamma delta T cells (**Supplementary Fig. 2**). Since the ICR transcriptional signature is largely reminiscent of T cell-driven immune responses, we investigated the association between T cell infiltration and the ICR score. Indeed, T cell infiltration was more abundant in ICR-high chordomas as compared to ICR-low samples (**Supplementary Fig. 3**). Of note, T cells were detected in all chordomas although the levels of infiltration were highly variable, both within and between lesions, ranging from 6.5 to 284.6 cells / mm^2^ (median = 51 cells / mm^2^) between tumours (**Fig. 2A, B**). Although the vast majority of T cells was confined to the stromal compartment surrounding the tumour lobules, most tumours also show infiltration of a low number of T cells within the cancer cell compartment. Other commonly observed immune cell populations included various subsets of myeloid cells, such as monocytes, macrophages, dendritic cells, and granulocytes (**Supplementary Fig. 2**).

**Fig. 2.**
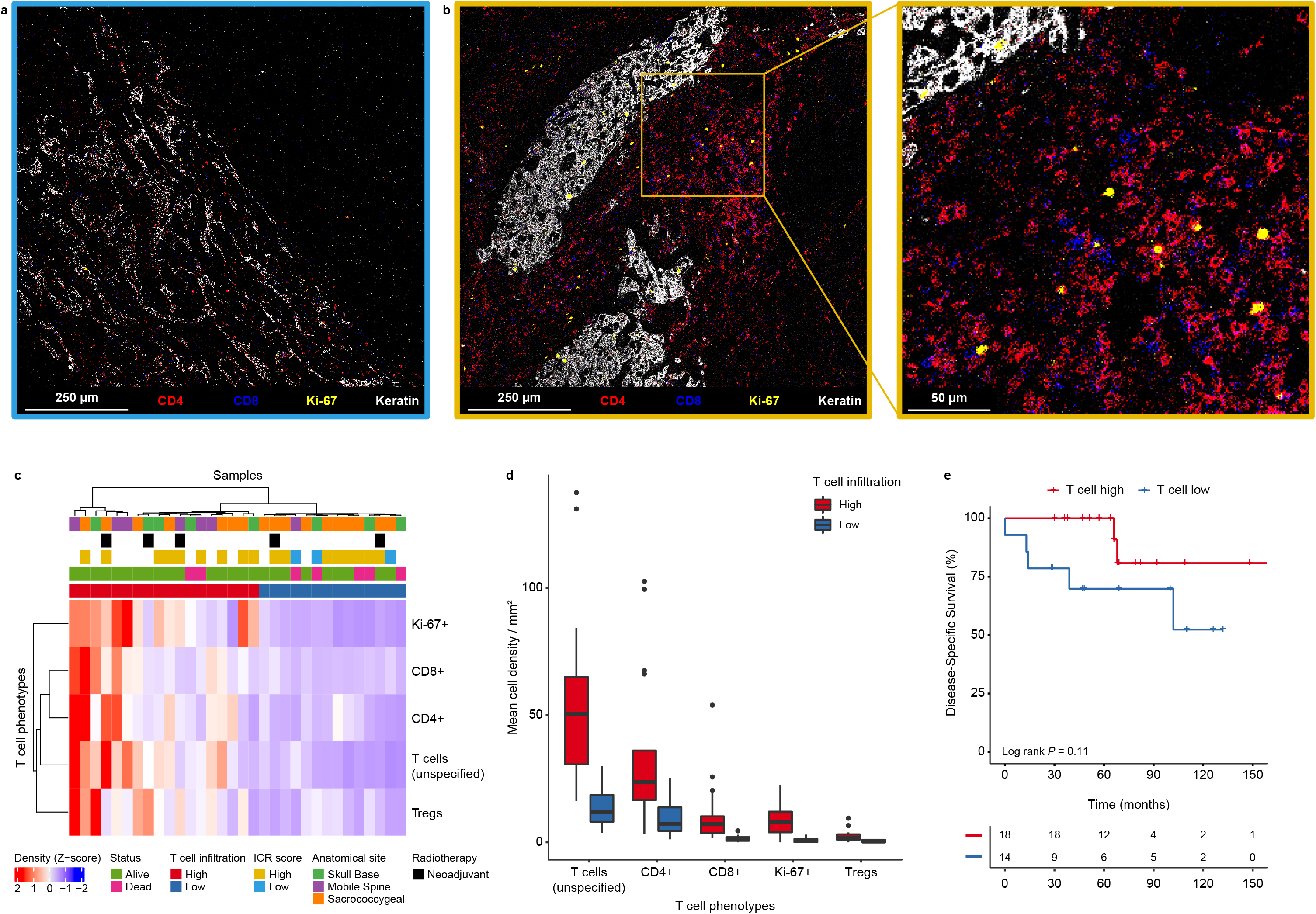
Chordomas present distinct groups based on T cell infiltration. **a-b**, Heterogeneity in T cell infiltration in chordomas. The images show T cell-related markers (CD4 = red, CD8 = blue), a proliferation marker (Ki-67 = yellow) and a tumour marker (keratin = white) as detected by imaging mass cytometry. **a**, chordoma with low T cell infiltration (ICR low). **b**, chordoma with high T cell infiltration and a magnified image of the chordoma with high T cell infiltration (ICR high). **c**, Heatmap showing the mean cell density Z-Score of all T cell phenotypes in all chordoma samples (*n* = 32), hierarchically clustered into a T cell high (*n* = 18) and T cell low (*n* = 14) group. Samples are annotated with their respective ICR score from the transcriptional analysis. In addition, the samples are annotated for their anatomical location, the follow-up status of the patients as well as whether the sample had received neoadjuvant radiotherapy. **d**, Boxplots presenting the mean cell densities per mm^2^ per classified T cell group (T cell high: *n* = 18; T cell low: *n* = 14) for the identified T cell phenotypes. **e**, Kaplan-Meier curves interrogating the association between T cell infiltration and disease-specific survival. Two-sided log-rank tests were performed for the Kaplan Meier estimates. ICR = immunologic constant of rejection; Tregs = regulatory T cells

Next, chordoma samples were clustered according to the extent of infiltration of the different T cell populations leading to the definition of two major groups (high vs. low T cell infiltration; **Fig. 2C-D**). All three chordoma samples previously classified as ICR low clustered in the T cell low group but samples that were previously classified as ICR high were equally distributed between high and low T cell groups (**Fig. 2C**), demonstrating the additional value of image-based evaluation of immune cell infiltration in solid tumours. The extent of T cell infiltration was not associated to whether a patient had received neoadjuvant radiotherapy, instead it was associated to the anatomical location of the tumours, with spinal chordomas enriched in the T cell high group and sacrococcygeal chordomas enriched in the T cell low group (**Fig. 2C**). Although it appeared that patients with higher T cell infiltration have an improved disease-specific survival, the observed association was not statistically significant (Log rank *P* = 0.11, **Fig. 2E**). There was no association between T cell infiltration and recurrence- or metastasis-free survival (**Supplementary Fig. 4A-B**). Moreover, unsupervised clustering of sequential tumours of six patients revealed that the immune contexture of chordomas can change upon disease recurrence or metastasis (**Supplementary Fig. 4C**).

### Antigen-presenting myeloid cells are abundant in T cell high chordomas and co-localise with T cells

Further exploration of the imaging mass cytometry data revealed that high T cell infiltration was accompanied by the presence of dendritic cells (T cell high vs T cell low, *P* < 0.001, Benjamini-Hochberg corrected false discovery rate [FDR] = 0.0052, **Fig 3A**). In addition, we found that T cell high chordomas were characterised by increased numbers of immune cell aggregates mainly composed of T cells and myeloid cells (*P* < 0.001, FDR = 0.0052, **Fig 3A**). Other phenotypes that were significantly associated with T cell infiltration include CD11c^+^HLA-DR^+^ macrophages (*P* = 0.014, FDR = 0.048), CD163^+^HLA-DR^+^ macrophages (*P* = 0.0079, FDR = 0.035), granulocytes (*P* = 0.0082, FDR = 0.035), podoplanin^+^ stromal cells (*P* < 0.001, FDR = 0.0052) and vessels (*P* = 0.0088, FDR = 0.035; **Supplementary Fig. 5A-B**). No specific phenotypes were associated with low T cell infiltration. Instead, this group exhibited a significantly lower density of immune cells (*P* < 0.001) and stromal cells (*P* < 0.01), resulting in a higher relative proportion of tumour cells (**Supplementary Fig. 5C**).

**Fig. 3.**
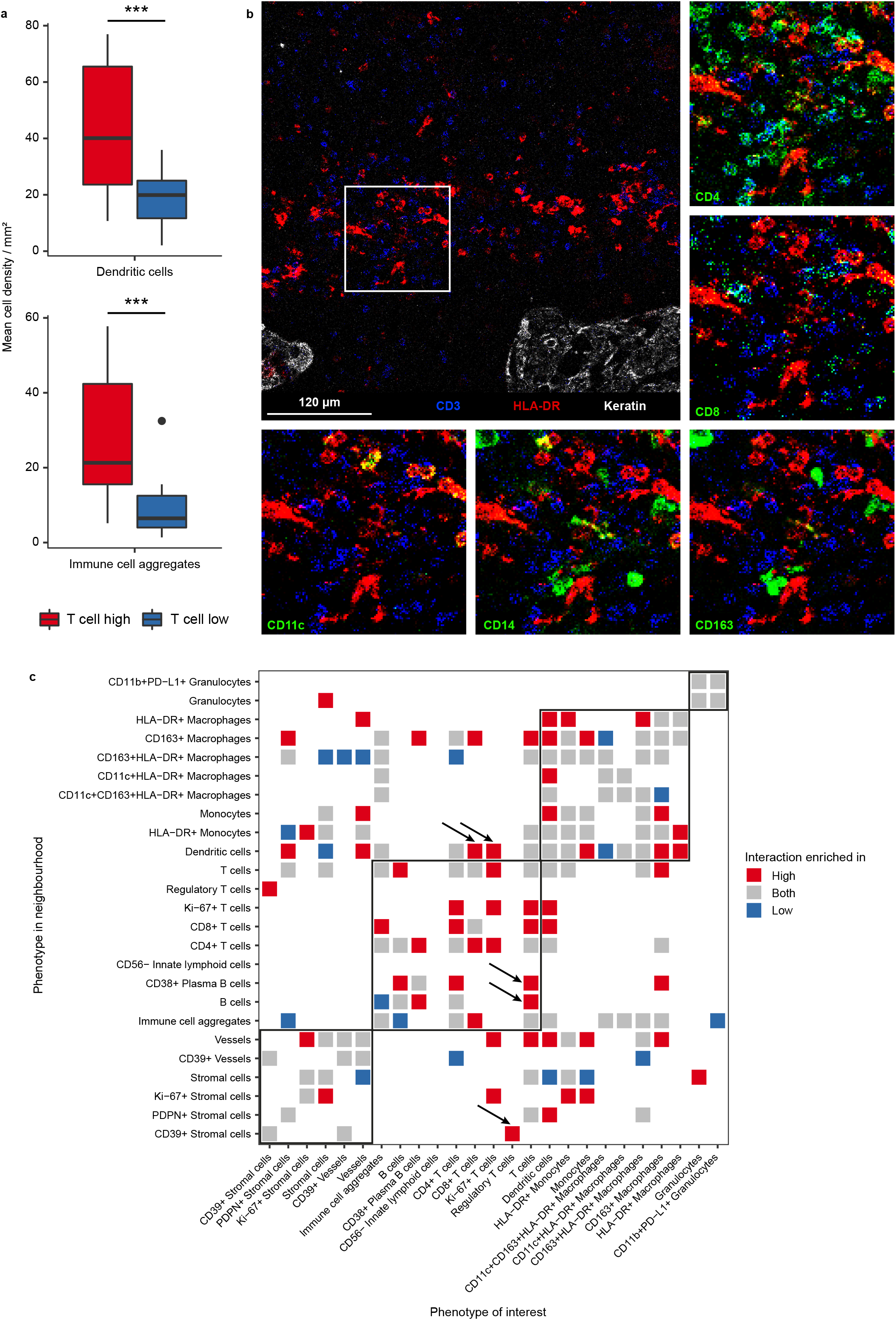
Pro-inflammatory interactions shape the immune microenvironment of T cell high chordomas. **a**, Boxplots comparing cell densities between the T cell high (*n* = 18) and T cell low groups (*n* = 14). Dendritic cells (CD11c^+/-^HLA-DR^+^CD68^-^) and immune cell aggregates (CD45^+^ and multiple differentiation markers, e.g. CD3^+^CD68^+^) are significantly higher in the T cell high group as compared to the T cell low group. Two-sided t-test was performed in R and the *P* value was adjusted with the Benjamini-Hochberg false discovery rate (FDR). * = *P* < 0.05; ** = *P* < 0.01; *** *P* < 0.001. **b**, An image generated with imaging mass cytometry displaying co-localisation of dendritic cells (HLA-DR = red) and T cells (CD3 = blue). Tumour cells are only shown in the overview image (top-left image, keratin = white). Other myeloid and T cell-related markers are presented in separate images around the overview image in green (CD4, CD8, CD11c, CD14 and CD163), to show co-localisation of different markers. **c**, Heatmap displaying the spatial interaction analysis results comparing the microenvironment of T cell high with T cell low chordomas. Only significant interactions are visualised and coloured for the subgroup(s) in which they are enriched. Permutation Z scores below 10 were considered not significant and thus removed for visualisation. Interactions of interest are indicated by black arrows. PDPN = podoplanin.

A comparative neighbourhood analysis on the imaging mass cytometry data was conducted to reveal distinct interaction patterns between both T cell groups independent of their overall abundance (**Fig 3C**). Naturally, most interactions involving T cells were enriched in the T cell high group, where most T cell phenotypes interacted with each other, with the exception of regulatory T cells which were only enriched in the neighbourhood of CD39^+^ stromal cells (**Fig. 3C**) The pro-inflammatory nature of the microenvironment of T cell high chordomas was further reinforced by the observed interactions between dendritic cells and CD8^+^ and Ki-67^+^ T cells as well as CD38^+^ plasma B cells and CD4^+^ and unspecified T cells (**Fig. 3C**). Most notably, the co-localisation of Ki-67^+^ T cells with CD4^+^ T cells and dendritic cells suggests interaction between these anti-tumour immune cells in a coordinated manner. In contrast, the most dominant interactions in the microenvironment of T cell low chordomas involved macrophage and stromal cell subsets including CD163^+^HLA-DR^+^ macrophages, stromal cells, vessels and CD39^+^ vessels (**Fig. 3C**). These observations suggest that the presence of T cells in T cell high chordomas accompanies a coordinated inflammatory response that also involves antigen-presenting myeloid cells such as dendritic cells, whereas the T cell low chordomas appear to be dominated with myeloid and stromal cell interactions.

### The extent of T cell infiltration in chordomas is independent of clinical parameters

To further explore the relation between the microenvironment of chordomas and the clinical behaviour of the disease, we performed multispectral immunofluorescence on the complete patient cohort (*n* = 76). A T cell-orientated antibody panel was designed to evaluate the expression of CD3, CD8, FoxP3, Ki-67 and pan-cytokeratin. Following cell phenotyping and automated counting, patients were clustered into T cell high and low groups (**Fig. 4A, B**). No significant association was found between T cell infiltration and disease-specific survival (Log-rank *P* = 0.17, **Fig. 4C**). Also, clinical features, such as anatomical location of the tumour or neoadjuvant radiotherapy, were not associated with T cell infiltration, suggesting that the presence of a coordinated immune response is not dependent on neoadjuvant radiation or the tumour’s location (**Fig. 4A**). Further associations between clinical parameters and prognosis were explored through cox proportional hazard models for the disease-specific, recurrence-free and metastasis-free survival, which are all presented in **Supplementary Table 3**. In short, previously established clinical parameters such as anatomical location of the primary tumour or the surgical resection margin were confirmed as prognostic factors.

**Fig. 4.**
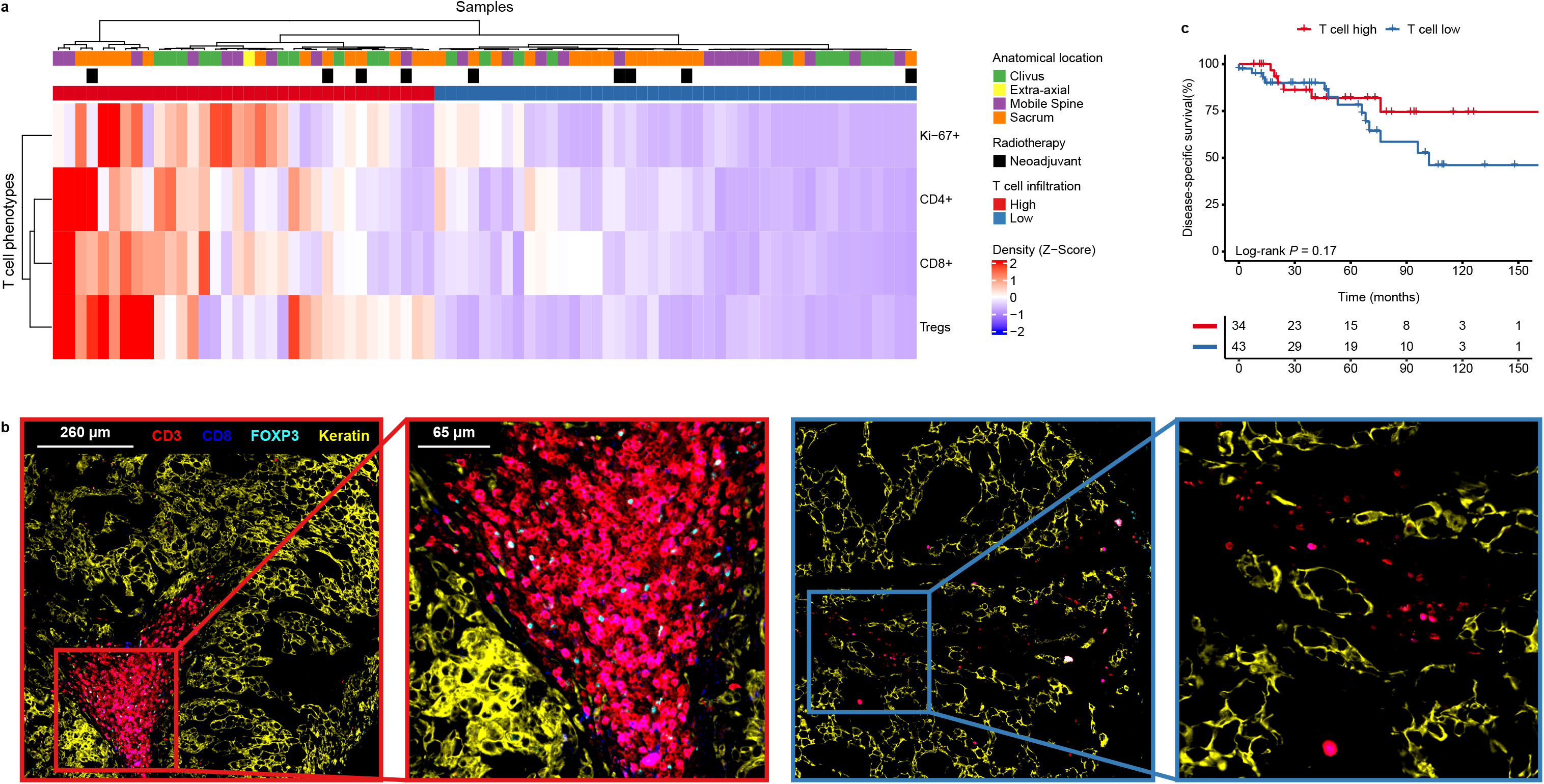
T cell intiltration is independent from clinical parameters. **a**, Heatmap displaying the mean cell density Z-Score per T cell phenotype as identitied with multispectral immunofluorescence in the entire cohort (*n* = 76). The samples are clustered in an unsupervised manner and thereby marked as T cell high (*n* = 34) or T cell low (*n* = 42). The samples are annotated for their respective anatomical location and whether they had received radiotherapy prior. **b**, Images displaying different T cell intiltration patterns as generated by multispectral immunofluorescence. The original and magnitied images show T cell-related markers (CD3 = red, CD8 = blue, FOXP3 = cyan) and a tumour marker (keratin = yellow) in two chordomas, each annotated for their respective T cell group (T cell high = red rectangles; T cell low = blue rectangles). **c**, Kaplan-Meier curves of the disease-specitic survival comparing the T cell high an T cell low groups of the entire cohort. Two-sided log-rank tests were performed for the Kaplan Meier estimates. Tregs = regulatory T cells.

### HLA class I expression is maintained in chordomas, but strong clonal expansion of T cells is rare

Strong immune selective pressure by T cells can lead to immune evasion by tumour cells, most notoriously through loss of HLA class I expression. Since high expression of HLA class I-related genes on RNA level does not preclude the loss of HLA class I expression due to, for instance, mutations in the *B2M* gene, we investigated the prevalence of HLA class I defects by immunodetection (**Fig. 5A**). In the complete cohort, HLA class I and β-2m membranous expression was observed in the majority of chordomas (73%). However, 22% of chordomas displayed defective HLA class I expression, which was accompanied by either loss or low expression of β-2m. The remaining 5% of the cases showed very weak expression of HLA class I and/or β-2m, which was identified by stronger stromal expression of HLA class I/β-2m but weak expression in the tumour cells. Thus, up to 27% of chordoma samples presented some form of impaired or altered antigen presentation. Interestingly, patients in the T cell low group more often had defective or weak HLA class I expression in comparison to T cell high patients (**Fig. 5B**). Positivity for HLA class I was relatively more frequent in T cell high patients (**Fig. 5B**). Intriguingly, we observed intratumoral γδ T cell infiltration in one T cell high chordoma lacking HLA class I expression (**Supplementary Fig. 6A-B**). γδ T cells have recently been proposed as effectors of immunotherapy in HLA class I-defective cancers^26^. These results indicate that HLA class I expression and, therefore, antigen presentation is maintained in the majority of chordomas.

**Fig. 5.**
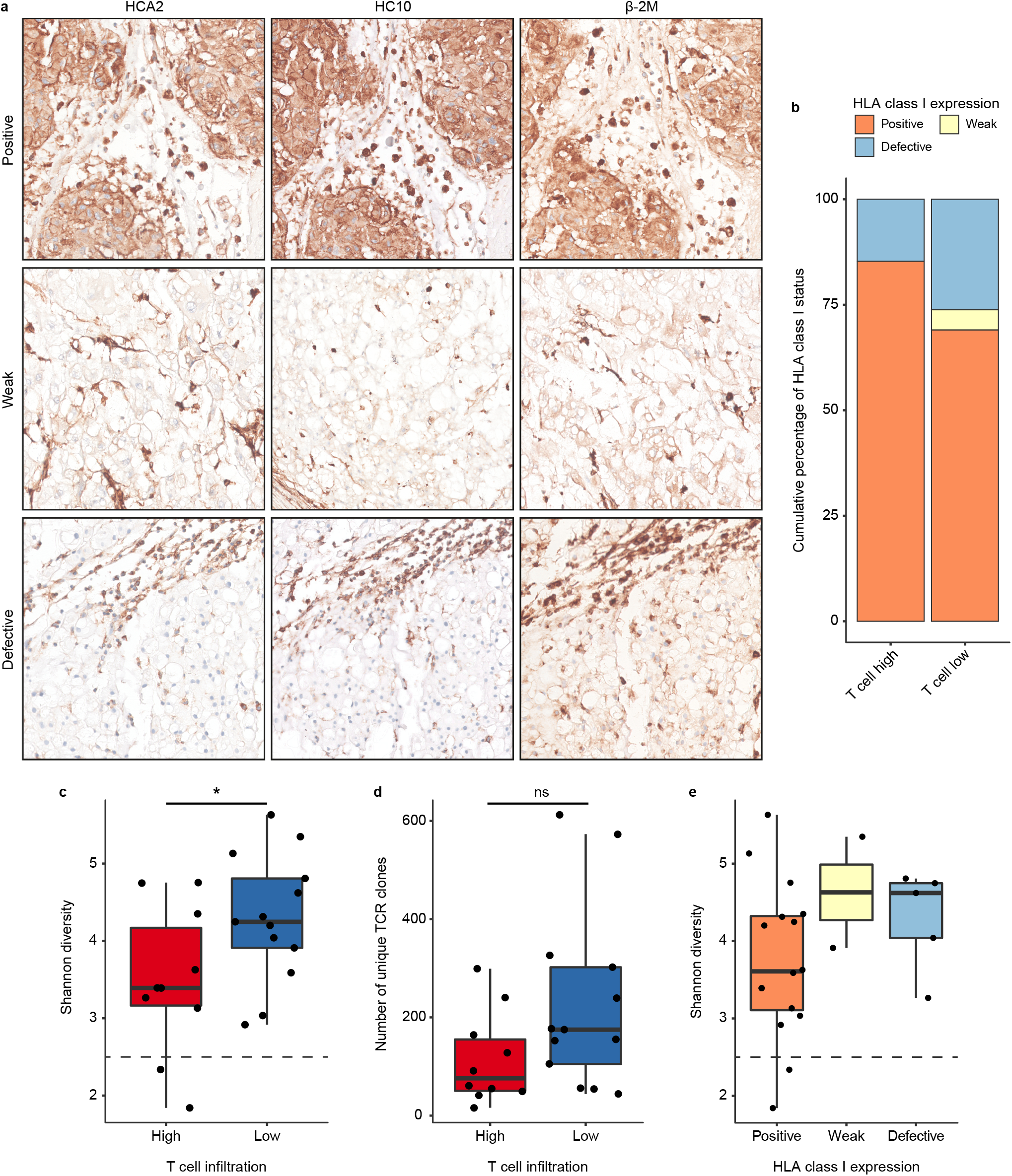
Antigen presentation and T cell clonal enrichment are present in chordomas. **a**, Human Leukocyte Antigen (HLA) class I (HCA2 and HC10) and β-2m expression patterns as validated by immunohistochemistry presenting the distinct expression patterns. From top to bottom, these images display positive expression of HLA class I and thus also β-2m, weak expression of HLA class I and β-2m and defective expression of HLA class I accompanied by weak expression of β-2m. **b**, Stacked bar plots displaying the percentual distribution of the observed HLA class I (both HCA2 and HC10) expression patterns in the chordoma samples in both T cell groups (T cell high: *n* = 34; T cell low: *n* = 40) **c-d**, Boxplots displaying the Shannon diversity (**c**) and the number of unique clones (**d**) of the profiled T cell receptors (TCR) from the immune microenvironment of chordomas (*n* = 23). Both the Shannon diversity and the number of unique clones are compared between the T cell high (*n* = 10) and the T cell low (*n* = 13) groups. Two-sided t-tests were performed to compare the T cell high and the T cell low group. * = *P* < 0.05; ns = not significant (*P* > 0.05). **d**, Boxplots presenting the Shannon diversity in relation to the observed HLA class I expression patterns in the subset that underwent TCR protiling (*n* = 22).

Presentation of tumour antigens can lead to the specific enrichment of T cell clones with anti-tumour specificity in the tumour tissues. To investigate the clonality of T cell responses in chordomas, we sequenced the variable region of the *TCRB* locus in 27 chordoma samples from 23 patients. Intriguingly, T cell high chordomas presented a more focused TCR repertoire than T cell low chordomas, which was demonstrated by the lower number of unique clones (*P* = 0.068) and Shannon diversity (*P* = 0.047) in T cell high chordomas compared to T cell low chordomas (**Fig 5C-D**). Additionally, tumours displaying a Shannon diversity below 2.5 were only contained within the T cell high group, suggesting the presence of strong clonal expansion in two T cell high chordomas (**Fig. 5C**). To substantiate the evidence for enriched TCR clonality, the Shannon diversity was compared to the HLA class I expression pattern of these 23 tumours, which revealed that patients with positive HLA class I expression, presented a more enriched TCR repertoire than patients with either defective or weak HLA class I expression (**Fig. 5E**).

## Discussion

Through a multimodal approach comprising RNA sequencing, imaging mass cytometry, multispectral immunofluorescence and TCR profiling, we studied the immune microenvironment of in total 77 chordoma patients. We found that chordomas present many hallmarks of anti-tumour immunity. We observed the majority of chordomas to have an “immune hot” microenvironment based on their immune-related transcriptomic profile, especially compared to other bone sarcomas (e.g. chondrosarcoma and osteosarcoma). Spatial analysis of immune cell infiltration further defined groups of chordoma patients based on the prevalence of T cell phenotypes. Increased T cell infiltration was associated with an apparently coordinated anti-tumour immune response as indicated by the presence of dendritic cells and other immune cells in immune cell aggregates. In contrast, chordomas with lower T cell infiltration were characterised by a microenvironment dominated by interactions comprising stromal cells and myeloid cells. Neither group was associated with clinical outcome nor with any other clinical parameter.

Recently, several studies have contributed to the improved understanding of the chordoma microenvironment, based on approaches including immunohistochemistry, multispectral immunofluorescence or single cell-RNA sequencing^6,7,23,27,28^. These studies identified myeloid cells and T cells as the most predominant immune cell populations in chordomas^7,23,27,28^. Similar to our findings, the majority of these immune cells was confined to the stroma, although both cell types could be found in the tumour nodules^6,7,28^. Moreover, other lymphoid cells such as NK cells (or innate lymphoid cells) and B cells were considered rare in chordomas^7,23,28^. Using imaging mass cytometry, we expanded on the previously identified immune cell populations and were able to establish immunologically relevant interactions within the chordoma microenvironment.

Studies focussed on methylation and RNA sequencing have identified groups of patients with either an inflamed or non-inflamed phenotype and associated these groups to clinical outcome, all with varying implications^29–31^. For instance, Zuccato *et al.* found through methylation profiling that a group of patients presenting an inflamed phenotype had a worse prognosis than patients without these inflammatory features^30^, whereas Huo *et al.* observed that patients from the non-inflamed group had a worse clinical outcome^29^. Furthermore, Baluszcek *et al.* described two similar groups based on methylation and transcriptomic profiling, without finding an association between these groups and survival^31^. All these studies, however, either only include skull base chordomas or skull base chordomas combined with spinal chordomas. The variable composition of the cohorts based on institutional referral patterns complicates the identification of common prognostic markers across studies. Here we included tumours from all major anatomical locations as well as different treatment strategies, providing an extensive characterisation of a uniquely large series of chordomas as representative of the general patient population as possible.

Nevertheless, most studies, including ours, support the separation of distinct groups of chordomas based on the abundance of pro-inflammatory immune cells in their microenvironment. Little is known, however, regarding potential drivers of these different immune contextures. In other tumour types (e.g., melanoma, carcinomas) spontaneous immune responses can often be explained by the mutation burden of a cancer where tumours with high mutation burden more often experience spontaneous anti-tumour immune responses as a consequence of high neoantigen abundance. In contrast, similar to most other sarcomas, the number of somatic, coding mutations in chordomas is generally low and stable between patients making it unlikely that distinct immune responses can be explained by the overall mutation burden of the tumours^13–15,32^. Although we only found evidence for strong clonal expansion in two T cell high chordomas, the TCR repertoire was less diverse in T cell high chordoma than T cell low chordomas, which would support that antigen recognition is a potential driver of immunity in chordomas. Naturally, this raises the question whether there is some specificity for the known chordoma-related transcription factor brachyury (*TBXT*) in these tumours, as it has increasingly been demonstrated that it can be recognised by T cells as well as that it can be targeted by means of vaccination in chordomas^9,33^. Another possible underlying factor to these distinct groups could be the extent of immunosuppressive mechanisms in the chordoma microenvironment. For instance, we identified abundant CD39 expression on vessels and stromal cells, in the vicinity of immune cells in both T cell high and T cell low chordomas, suggesting that the adenosine pathway might be a potential mediator of immune suppression in chordomas. Although the immunosuppressive functionalities of this pathway have mainly been described in carcinomas^34,35^, expression of CD39 was recently found to associate with increased inflammation in primary tumours as well as with increased T cell densities and PD-1 expression in recurrent undifferentiated soft tissue sarcomas^36^. Further research should determine the extent to which the adenosine pathway can modulate immune responses in this disease.

Going forward, it will be crucial to understand how the chordoma microenvironment impacts the outcomes of immunotherapy, in particular immune checkpoint blockade therapy. Intuitively, and as observed in other tumour types, responses should be more likely to occur in patients where spontaneous anti-tumour immune responses are observed. Encouragingly, our data also supports the notion that most chordoma patients may be eligible to immunotherapeutic approaches as: 1) despite differences in the extent of immune infiltration between tumours, the majority of chordomas display enhanced hallmarks of anti-tumour immunity as compared to other sarcoma types and 2) we determined that the majority of chordoma cases retain HLA class I expression and, therefore, antigen presentation. Therefore, other approaches including antigen vaccination or T cell based immunotherapies are, in theory, applicable for chordoma patients and may be particularly important in chordomas that do not present with a conspicuous anti-tumour immune response. We speculate that providing immunotherapy in the neoadjuvant setting^37^ to chordoma patients, either with or without radiotherapy, might have a strong impact on the clinical management of these tumours as it might improve surgical outcomes through tumour shrinkage. This could contribute to less extensive and complicated surgical procedures which could either reduce the risk of surgery-related morbidities (e.g. by saving critical nerve structures in the axial skeleton) or possibly facilitate complete surgical resection.

In conclusion, through a multimodal approach we provided evidence that chordomas present multiple hallmarks of anti-tumour immunity. On a transcriptional level, chordomas appear more enriched for inflammatory signalling than other genetically complex sarcomas. Within our chordoma cohort, we identified distinct immune contextures, either defined by a coordinated, possibly antigen-driven, immune response or a microenvironment dominated by stromal- and myeloid cell interactions. Furthermore, antigen presentation was maintained in the majority of chordomas. Our study provides a rationale for utilising immunotherapeutic strategies to aid in the clinical management of chordomas.

## Methods

### Patient samples

Formalin-fixed paraffin embedded (FFPE) and snap-frozen tissue material was initially collected for 43 and 24 chordomas respectively from 32 patients, followed by a second expansion cohort containing FFPE material of an additional 53 patients. A waiver of consent was obtained from the medical ethical evaluation committee (protocol number: B17.039 and B21.009). All specimens in this study were pseudo anonymised and handled according to the ethical guidelines described in ‘Code for Proper Secondary Use of Human Tissue in The Netherlands’ of the Dutch Federation of Medical Scientific Societies. Regions of interest (ROIs) representative for the tumour’s immune microenvironment were selected on Haematoxylin & Eosin (H&E) sections of decalcified FFPE tissue by a bone and soft tissue tumour pathologist (JVMGB). These ROIs were then used to create tissue micro arrays (TMAs) using a TMA Master (3DHISTECH). These TMAs were holding 2-4 cores of 1.6 mm in diameter per tumour, dependent on the source of the material or the morphology of the tumour (two cores: biopsy; three cores: resection/excision etc; four cores: heterogeneous morphology, or separate areas of dedifferentiation). As controls and for orientation purposes, each TMA contained three cores tonsil and three cores placenta, either decalcified with EDTA or formic acid or non-decalcified.

### RNA sequencing data

RNA was isolated from snap-frozen tissue from 20 primary tumours with TRIzol and Isopropanol/Ethanol. In addition, RNA sequencing data from primary tumours of osteosarcoma patients (*n* = 7; <u>GSE237033</u>), chondrosarcoma patients (*n* = 9), undifferentiated soft tissue sarcoma patients (*n* = 13), and myxofibrosarcoma patients (*n* = 10) was taken along as a means of comparison. RNA purification was performed with the RNeasy kit which included treatment with DNase. Paired-end 150 base pair (bp) reads were created on a NovaSeq6000 Illumina at GenomeScan (Leiden, The Netherlands). The sequencing data was processed with the RNA-seq BioWDL pipeline from the SASC with default settings (<u>BioWDL Github</u>, LUMC, The Netherlands). Alignment of the 150bp reads to hg38 was performed with STAR, after which the gene expression was quantified with HTSeq-count^38^. The processed sequencing data was stored as a matrix containing the counts per gene per sample.

### Gene expression analysis

Normalisation and exploratory analysis of the RNA sequencing data was performed in R (v.4.0.2) by using DESeq2 (v. 1.30.1). BiomaRt (R, v.2.46.3) was used to translate the ensemble IDs to known protein coding genes. The most variably expressed genes were determined by calculating the variance per gene across all samples. The immunologic constant of rejection (ICR) gene signature was used to describe the activity state immune microenvironment and to identify distinct immune-related clusters (Th1-related markers: *IFNG*, *STAT1*, *IL12B*, *IRF1*, *TBX21*; CD8+ T cells: *CD8A*, *CD8B*; Immune effector molecules: *GZMA*, *GZMB*, *GZMH*, *PRF1*, *GNLY*; Chemokine ligands: *CXCL9*, *CXCL10*, *CCL5*; Immune suppressive molecules: *IDO1*, *PDCD1*, *CD274*, *CTLA4*, *FOXP3*)^24^. The mean of the log2 normalised expression of these 20 genes was used to attribute an ICR “score” to each sample for comparison between the sarcoma subtypes, which was visualised with ggplot2 (R, v.3.3.6). For both the heatmaps in **Fig 1**, a Z-Score was calculated per gene to visualise the transcriptional profile of the samples by using the Ward variance method with ComplexHeatmap (R, v.2.11.1). Differential gene expression analysis was performed with DESeq2 comparing chordomas to all other sarcomas. Genes were considered differentially expressed when the log2 fold change was above 2 and the adjusted *P* value was below 0.01. Immune-related genes were selected for visualisation in the enhanced volcano plot and additional boxplots. For visualisation purposes, the axes of the enhanced volcano plot were manually adjusted as follows: log2 fold change (x-axis) was set between −10 and 10, and the −log10 adjusted *P* (y-axis) was set to a maximum of 100.

### Imaging mass cytometry

Conjugation of BSA-free antibodies and immunodetection with a 40-marker tumour immune microenvironment panel (**Supplementary Table 4**) was performed as previously described^39^. In brief, four-µm sections of the created TMAs were deparaffinised and rehydrated, after which heat-induced antigen retrieval was performed in 1x low pH antigen retrieval solution (pH6, Thermo Fisher Scientific). Subsequently, the first batch of primary antibodies were incubated overnight at 4 °C. After washing in PBS supplemented with 1% BSA and 0.05% Tween, conjugated anti-mouse and anti-rabbit antibodies were incubated for one hour at room temperature. Next, the sections were incubated with the second batch of primary antibodies for five hours at room temperature, followed by an overnight incubation at 4 °C with the final batch of primary antibodies. To finish, 1.25 µM of the DNA intercalator Iridium (Fluidigm) was incubated for five minutes to stain DNA, after which the sections were washed, air-dried and stored at room temperature. ROIs of 1000×1000 µm were selected with consecutive H&E slides and ablated by using a Hyperion mass cytometry imaging system (Fluidigm) at the Flow Core Facility (LUMC, Leiden) within a month after staining. The imaging data was acquired with CyTOF Software (v.7.0), visually inspected and then exported with MCD Viewer (v.1.0.5). In the end, 127 tumour ROIs were successfully ablated and analysed.

### Imaging mass cytometry analysis

To correct for variability in signal-to-noise ratios between FFPE sections, the signal intensity of the generated images was normalised at pixel levels by using Matlab (v.R2021a), after which the images were binarised using semi-automated background removal in Ilastik (v.1.3.3) as described previously ^39^. Probability masks were generated in Ilastik by using a multicoloured image including the DNA channel, CD45 channel, Keratin channel and Vimentin channel to annotate nucleus, cytoplasm and background. Probability masks were loaded into CellProfiler (v.2.2.0) and used for creating cell segmentation masks. Next, the segmentation masks and binarised images were loaded into ImaCyte^40^ to generate single cell-FCS files containing relative frequency of positive pixels for each marker, which were then used for phenotyping in CytoSplore (v.2.3.1) by sequentially utilizing Hierarchical Stochastic Neighbour Embedding (H-SNE) and t-distributed Stochastic Neighbour Embedding (t-SNE). Identified cell phenotypes were loaded into ImaCyte together with the segmentation masks and binarised images, which allows the mapping of the phenotypes back onto the images. Phenotypes per image were extracted to calculate the mean cell density per mm^2^ for every sample (one image = one mm^2^). An overview of the lineage markers used for phenotype identification is listed in **Supplementary Table 5**. Patients were clustered into two groups based on the Z-Score normalised mean cell densities per mm^2^ of all five identified T cell phenotypes by using the Ward variance method with ComplexHeatmap. Ggplot2 was used to visualise the mean cell densities and to compare them between both T cell groups. For statistical testing, a two-sided student’s t-test was performed followed by a Benjamini-Hochberg FDR correction. In addition, an overview heatmap containing all identified phenotypes in the microenvironment of chordomas was visualised with ComplexHeatmap.

### Spatial interaction analysis

After phenotype calling, a spatial interaction analysis was performed with ImaCyte for all imaged samples, comparing T cell low with T cell high chordomas. ImaCyte utilises a 1000-iteration permutation test which calculates a Z-Score for the interaction between adjacent cells. However, permutation testing does not adjust for overabundant cell types, such as tumour cells, and is therefore not able to identify meaningful interactions for tumour cells. Because of the range of the calculated Z-Scores, interactions with a Z-Score below 10 were considered not significant and therefore removed for visualisation only. If an interaction Z-Score was higher than 10 in one group and lower than 10 in the other group, this interaction was considered significant for one group. When both Z-Scores were higher than 10, this interaction would be considered significant in both groups. A comparative interaction heatmap between all phenotypes excluding tumour cells was visualised with ggplot2.

### Multispectral immunofluorescent imaging of T cells

A T cell panel comprising five markers for five channels was generated for imaging with a Vectra (Akoya Biosciences) as previously described^41^ for validation purposes. This panel included FOXP3 (236A/E7, 1:1000, Invitrogen, 14-4777-82), CD8α (D8A8Y, 1:100, Cell Signaling Technology, #85336), CD3ε (EP449E, 1:50, Abcam, ab52959), pan-cytokeratin (AE-1/AE-3 & C11, both 1:50, Novus & Santa Cruz, NBP2-33200AF647 & SC-8018) and Ki-67 (8D5, 1:200, Cell Signaling Technology, #9449) and DAPI as a nuclear staining. In short, slides were deparaffinised and incubated in 0.3% H_2_O_2_ in methanol for 20 minutes to block endogenous peroxidase. After antigen retrieval in citrate (pH 6.0), slides were incubated for 30 minutes with SuperBlock^TM^ Blocking Buffer (ThermoFisher Scientific). Next, slides were incubated with primary antibody (FOXP3) for one hour at room temperature and subsequently with BrightVision DPVO-HRP (Immunologic) for 30 min at room temperature. Opal 520 reagent (Akoya Bioscience) was diluted in Opal amplification diluent and then added to the slides for one hour at room temperature, after which the slides were heated in citrate buffer (pH 6.0) for 15 min. The second round of primary antibodies (CD8α and Ki-67) were incubated overnight at room temperature and then stained with their respective fluorescent secondary antibodies (CF555, 1:400, Biotum and Alexa 680, 1:200, Invitrogen) for one hour at room temperature. Finally, the slides were stained with directly conjugated antibodies (CD3ε-Alexa 594 and pan-cytokeratins-Alexa 647) for 5 hours at room temperature and then mounted. All TMAs were imaged at 20x, after which the images were processed with inForm (v.2.4), which comprises background removal for each antibody by normalisation of the spectra, and analysed with QuPath (v.0.3.1). Object classification was performed by using the following classes: CD3^+^FOXP3^+^ = Regulatory T cells, CD3^+^CD8^+^ = CD8^+^ T cells, CD3^+^ = CD4^+^ T cells, CD3^+^Ki-67^+^ = Proliferating T cells, CD3-= Rest. Counts per image (one image = 1 mm x 1.3 mm) were normalised per patient to counts per mm^2^ tissue. Patients were divided into a T cell high and low cluster based on the unsupervised clustering of the Z-scores of all T cell phenotypes in R.

### Survival analysis

For all chordoma patients, Kaplan-Meier curves for the disease-specific survival and recurrence-free survival were generated with the survminer and survival packages (R, v.0.4.9 and v.3.1-12 respectively). The disease-specific survival was considered the survival of a patient until death due to the disease, whereas the recurrence-free survival was considered the time until a patient was diagnosed with a recurrence, thereby excluding patients who did not receive surgery. Since the cohort included many censored cases, the log-rank *P* value was used to estimate the significance of the associations. Known prognostic factors such as the anatomical tumour location excluding the only extra-axial patient, age and type of treatment, surgical margin excluding patients who did not receive surgery and disease recurrence were investigated for their prognostic value, as well as the newly identified T cell infiltration groups. Univariate and multivariate analyses were performed in R according to the cox proportional hazard model. Only clinical parameters that were found significant in the univariate analysis were taken along in the multivariate analysis.

### Immunohistochemical detection of HLA class I and β-2m

Immunohistochemistry (IHC) was performed for the HLA class I heavy chain (HCA2, 1:1000, Nordic Bio, MUB2036P; HC10, 1:3200, Nordic Bio, MUB2037P) and β-2m (EPR21752-214, 1:4000, Abcam, ab218230) on all generated TMAs as described previously^42^. Briefly, antigen retrieval was performed in citrate (pH 6.0), after which the TMA sections were pre-incubated with PBS-5% non-fat dry milk for 30 min at room temperature. Primary antibody dilutions were incubated overnight at 4 °C, after which the sections were incubated with BrightVision DPVO-HRP (Immunologic) for 30 min at room temperature. Finally, TMA sections were further developed with liquid DAB+ chromogen (Dako) for 10 min and subsequently counterstained with haematoxylin. Samples were classified as HLA class I positive, weak, or defective depending on the expression pattern of HC10 and HCA2 in the tissue and the contrast between the expression of the tumour cells and the immune cells. Tumours were classified as defective if one or more of the HLA class I antibodies were negative in the tumour cells. The distribution of the expression patterns among the identified T cell groups was visualised in R using ggplot2.

### TCR β-chain sequencing

DNA was isolated from snap-frozen tissue from 27 chordomas from 23 patients for sequencing of the variable region of the *TCRB* locus. In brief, DNA libraries were generated with the AmpliSeq Library Kit Plus and the OncoMine TCRβ-SR Assay following the manufacturer’s instructions. Sample libraries were measured with the ION Library TaqMan Quantitation Kit and then pooled to a final concentration of 25 pM. Next, the pooled libraries were templated on ION 540 Chips with the Ion Chef System. The 80bp reads were created on an Ion GeneStudio S5 Prime Sequencing Technology at GenomeScan (Leiden). The number of clones and the Shannon diversity, a measure for clonal enrichment, were calculated in R by using productive counts only, meaning that a clone needed at least 10 plus and 10 minus counts. Because recurrences showed similar levels of TCR diversity as their respective primary tumour, we used the TCR repertoire of recurrences for three patients for which there was no frozen material of primary tumours available. Finally, the data was visualised in R using ggplot2.

### Code & data availability

The RNA sequencing data generated in this study are publicly available on GEO (<u>GSE239531</u>). The imaging mass cytometry data is available on BioImage Archive (<u>S-BIAD830</u>). The RNA sequencing data for the other studied sarcoma subtypes is unpublished and therefore not yet available, with the exception of the osteosarcomas (<u>GSE237033</u>). The used code and processed data will be published on GitLab for reproducibility. All other data are available from the corresponding author on reasonable request.

## Acknowledgments

The authors would like to thank the Flow Core Facility and the Information Technology & Digital Innovation department of the Leiden University Medical Center for their service.

## Author contributions

**S.v.O.** contributed to the conceptualisation, methodology, investigation, data curation, data analysis, visualisation, writing-original draft and writing-review editing. **D.M.M.** contributed to the conceptualisation, investigation, data curation, data analysis and writing-review editing. **M.E.IJ.** contributed to the experiments, data analysis and writing-review editing. **J.R.** contributed to the data analysis and writing-review editing. **B.v.d.A.** contributed to the investigation and writing-review editing. **R.v.d.B.** contributed to the investigation and writing-review editing. **I.H.B.d.B.** contributed to the investigation and writing-review editing. **M.v.d.P.** contributed to the investigation and writing-review editing. **P.M.W.K.** contributed to the investigation and writing-review editing. **S.B.P.** contributed to the resources and writing-review editing. **W.C.P.** contributed to the resources and writing-review editing. **R.J.P.v.d.W.** contributed to the resources and writing-review editing. **N.F.C.C.d.M**. contributed to the conceptualisation, methodology, investigation, supervision, writing-original draft and writing-review editing. **J.V.M.G.B.** contributed to the conceptualisation, methodology, investigation, supervision, writing-original draft and writing-review editing.

## Index of supplementary information

**Supplementary Fig. 1** | Chordomas are enriched for immune-related gene signaling

**Supplementary Fig. 2** | Overview of the chordoma microenvironment

**Supplementary Fig. 3** | ICR high chordomas are enriched with T cell phenotypes

**Supplementary Fig. 4** | The immune contexture of chordomas varies over time

**Supplementary Fig. 5** | Comparative overview of the chordoma immune microenvironment

**Supplementary Fig. 6** | γδ T cell infiltration in relation to HLA class I expression

**Supplementary Table 1** legend (table added separately)

**Supplementary Table 2** legend (table added separately)

**Supplementary Table 3**. Univariate and multivariate cox proportional hazard results in the combined cohort

**Supplementary Table 4**. Imaging mass cytometry marker panel

**Supplementary Table 5**. Cell type lineage markers imaging mass cytometry

**Supplementary references**

**Supplementary Fig. 1.**
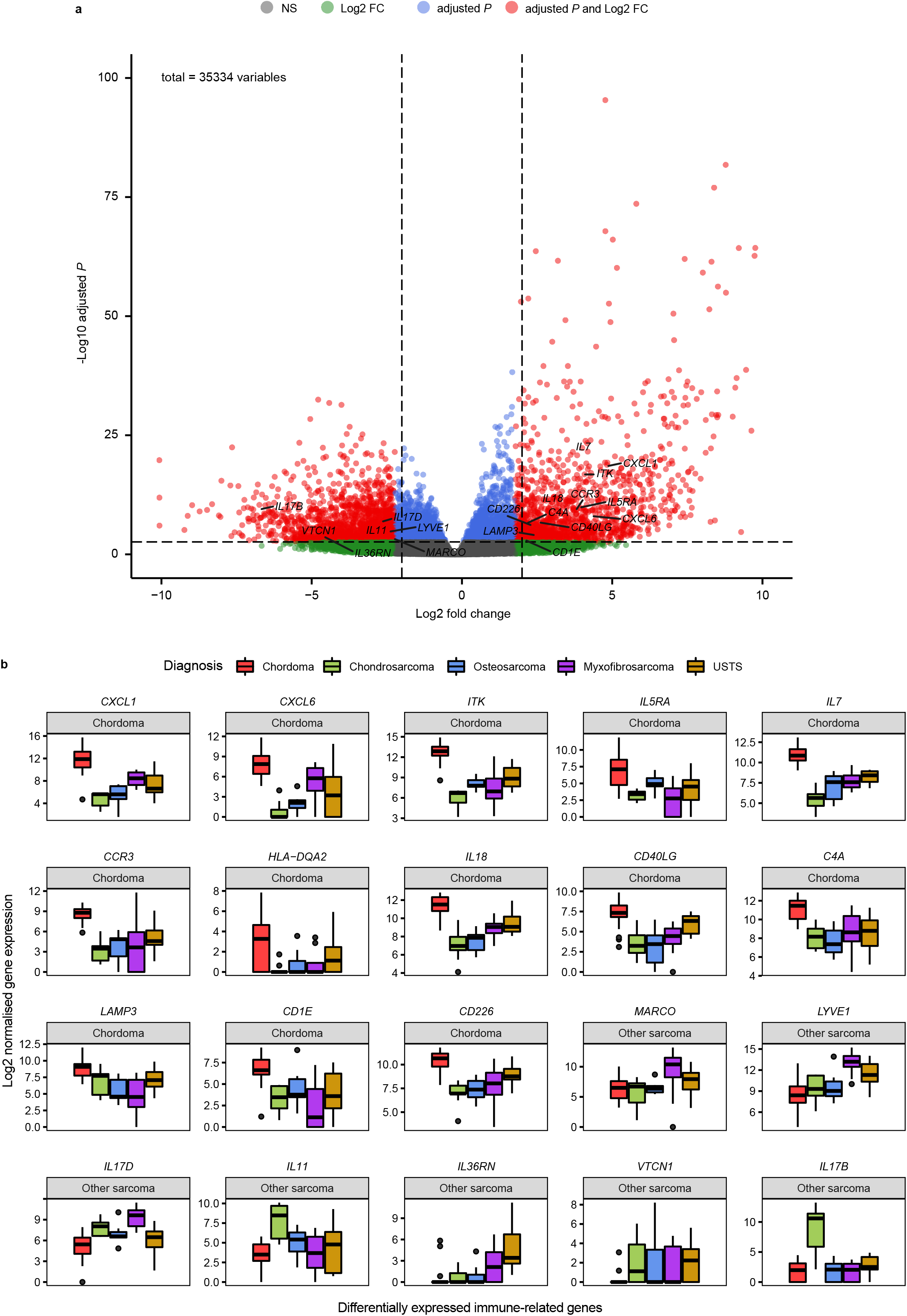
Chordomas are enriched for immune-related gene signaling. **a,** Enhanced volcano plot presenting the differential gene expression analysis between chordomas (*n* = 20) and all other studied sarcomas (*n* = 39. Genes with a log_2_ fold change above 2 and an adjusted *P* value below 0.01 were considered significantly differentially expressed. Immune-related genes were labelled for visualisation. **b**, Boxplots displaying the log_2_ normalised expression of the differentially expressed immune related genes. The boxplots are ordered in descending order for the log_2_ fold change and annotated for the respective group in which they are significanly enriched (chordoma vs other sarcoma). The boxplots are coloured for their respective histotype. NS = not significant; FC = fold change; USTS = undifferentiated soft tissue sarcoma.

**Supplementary Fig. 2.**
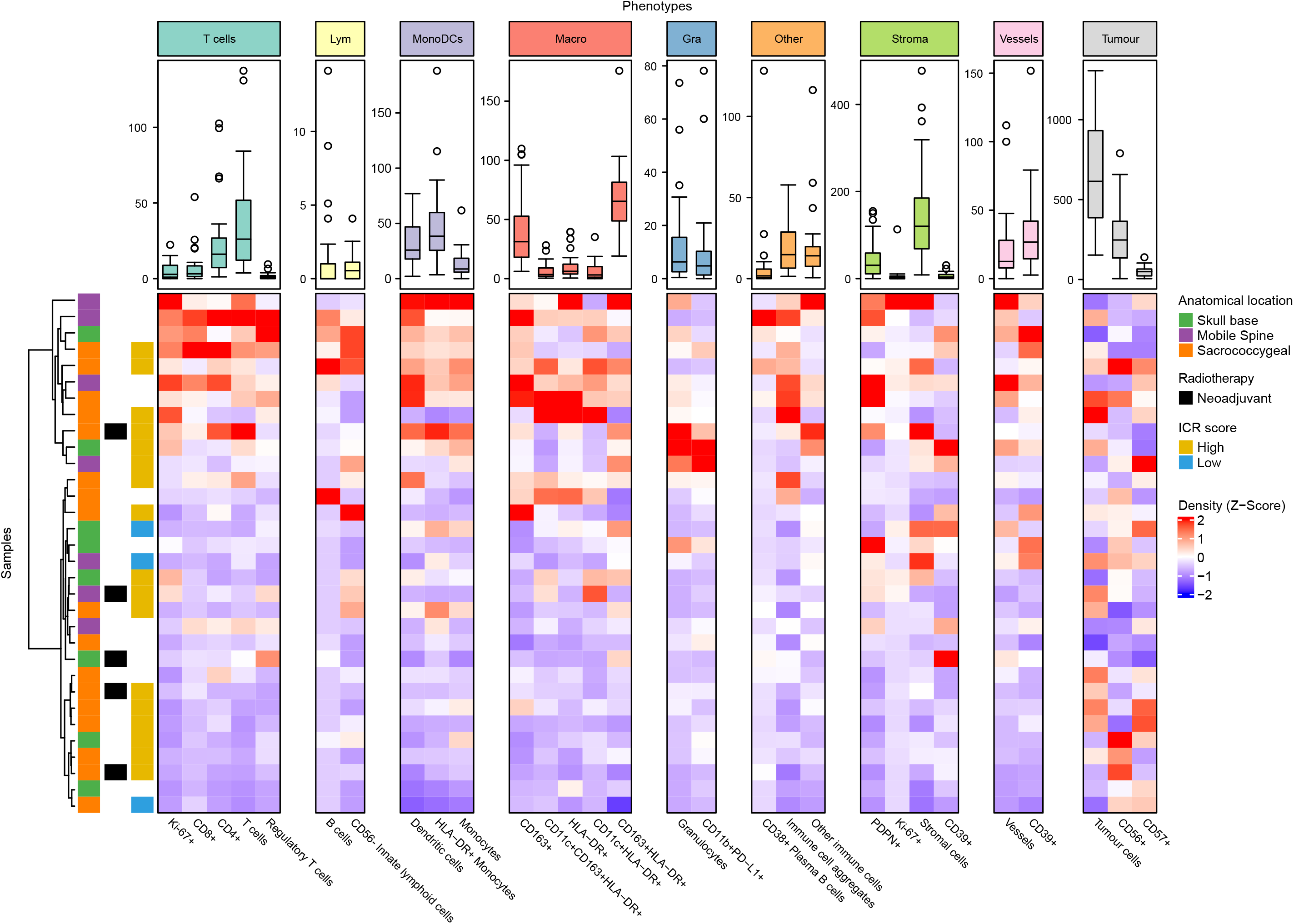
Overview of the chordoma microenvironment. Overview heatmap of the mean cell density Z-Score of 29 identitied cell phenotypes in all chordoma samples (*n* = 32). The samples that were used for transcriptome sequencing are annotated according to their ICR score. Samples are hierarchically clustered and annotated for their anatomical location and whether they had received neoadjuvant radiotherapy. Boxplots are presented per phenotype to display the general mean cell densities per mm^2^ for each phenotype. ICR = immunologic constant of rejection; Lym = lymphoid; MonoDCs = monocytes and dendritic cells; Macro = macrophages; Gra = granulocytes; PDPN = podoplanin.

**Supplementary Fig. 3.**
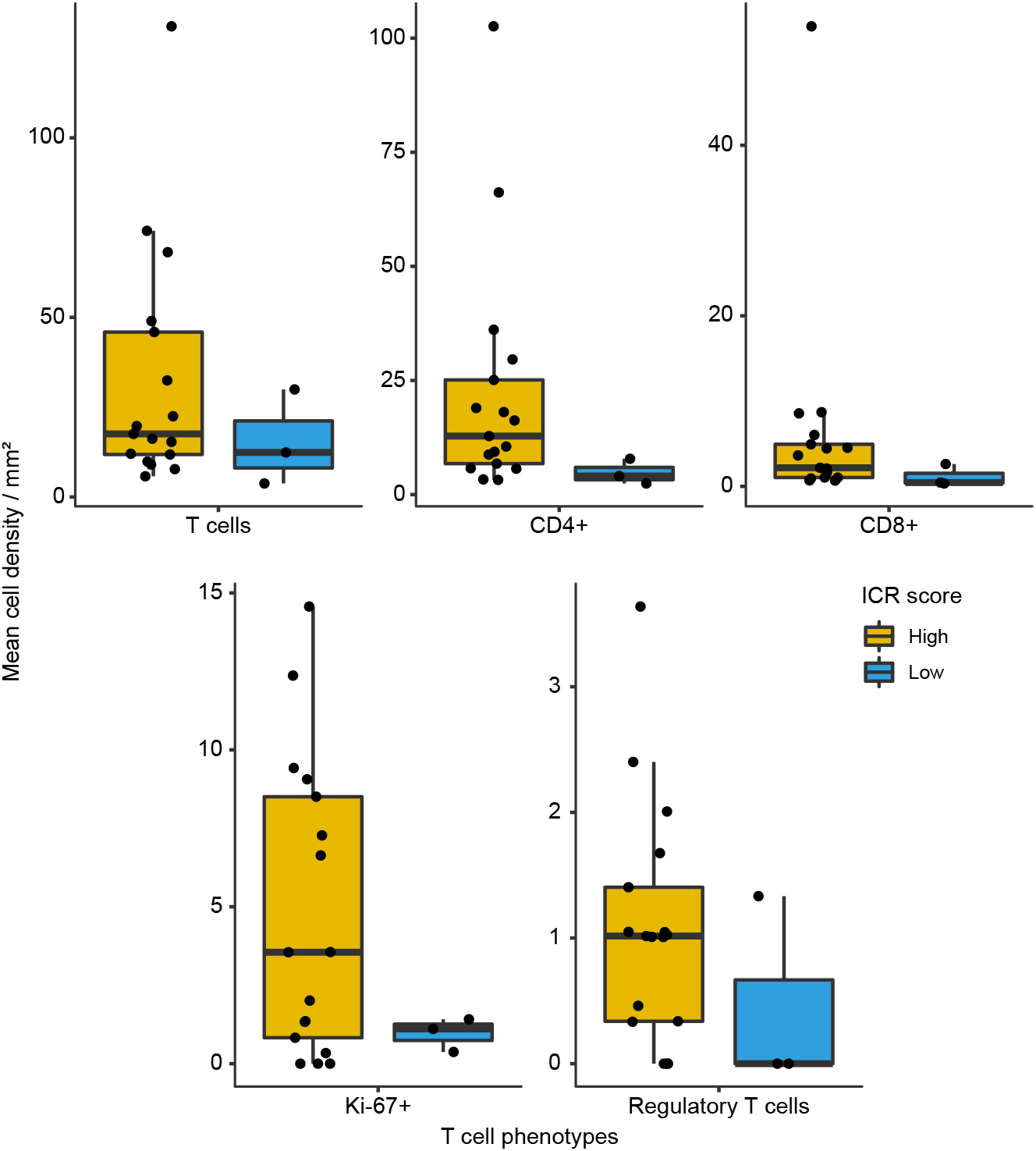
ICR-high chordomas are enriched with T cell phenotypes. Boxplots representing the different levels of intiltration of the identitied T cell populations between ICR-high (n = 17) and ICR-low (*n* = 3) chordomas. No statistical tests were performed due to the small sample size. ICR = immunologic constant of rejection.

**Supplementary Fig. 4.**
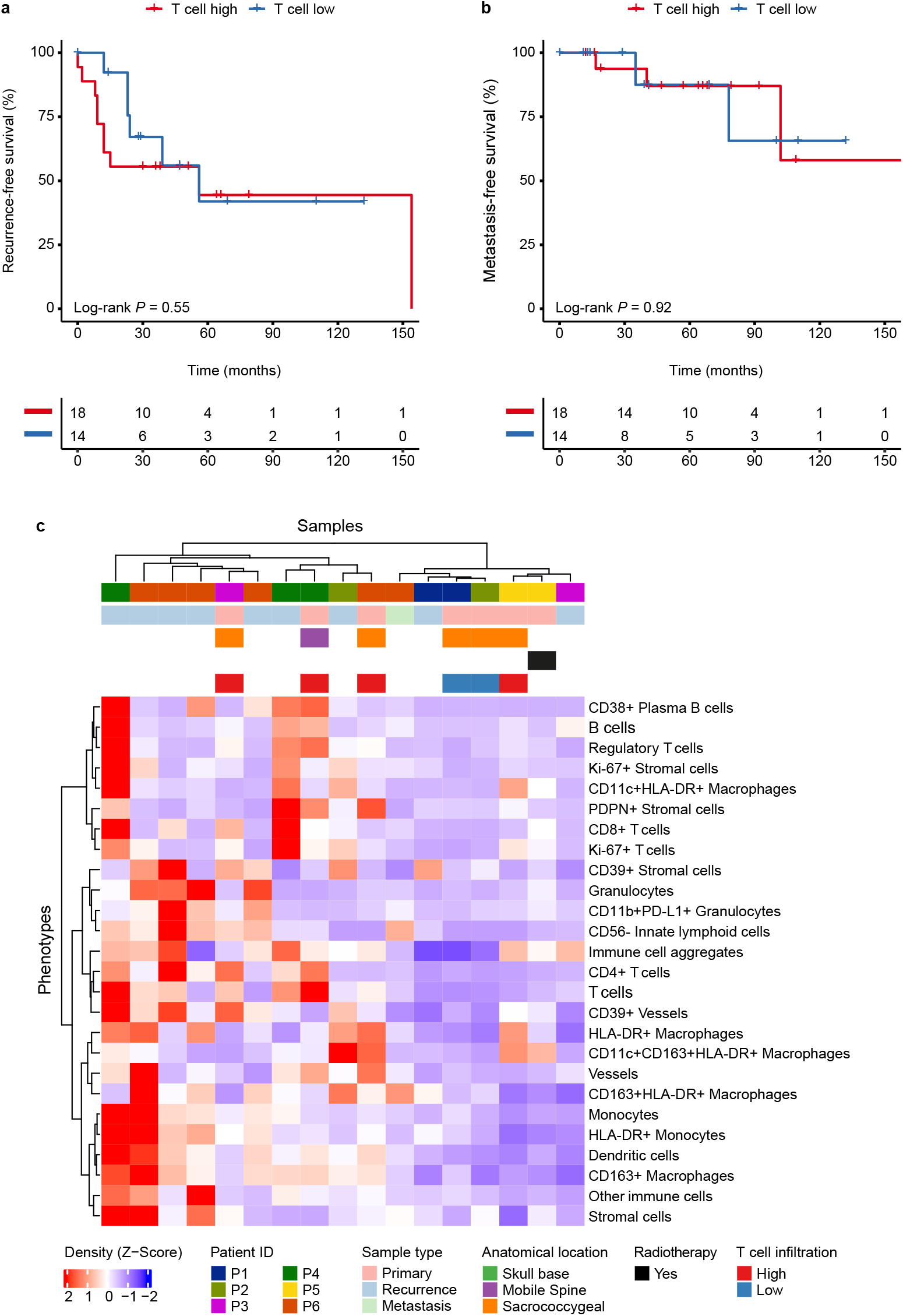
The immune contexture of chordomas varies over time. **a-b**, Kaplan-Meier curve indicating the association between T cell intiltration and recurrence-free survival (**b**) as well as metastasis-free survival (**c**). Two-sided log-rank tests were performed for the Kaplan Meier estimates. **c**, Heatmap presenting the unsupervised clustering of untreated primary tumours (*n* = 6) and their respective treated primary tumour (*n* = 1), recurrences (*n* = 9) and metastases (*n* = 1). The heatmap displays the mean cell density Z-Score of the identitied cell phenotypes, excluding tumour cells. The samples are annotated for patient ID, type of sample, anatomical location, whether the tumour was treated with neoadjuvant radiotherapy and T cell group of the primary tumour. PDPN = podoplanin.

**Supplementary Fig. 5.**
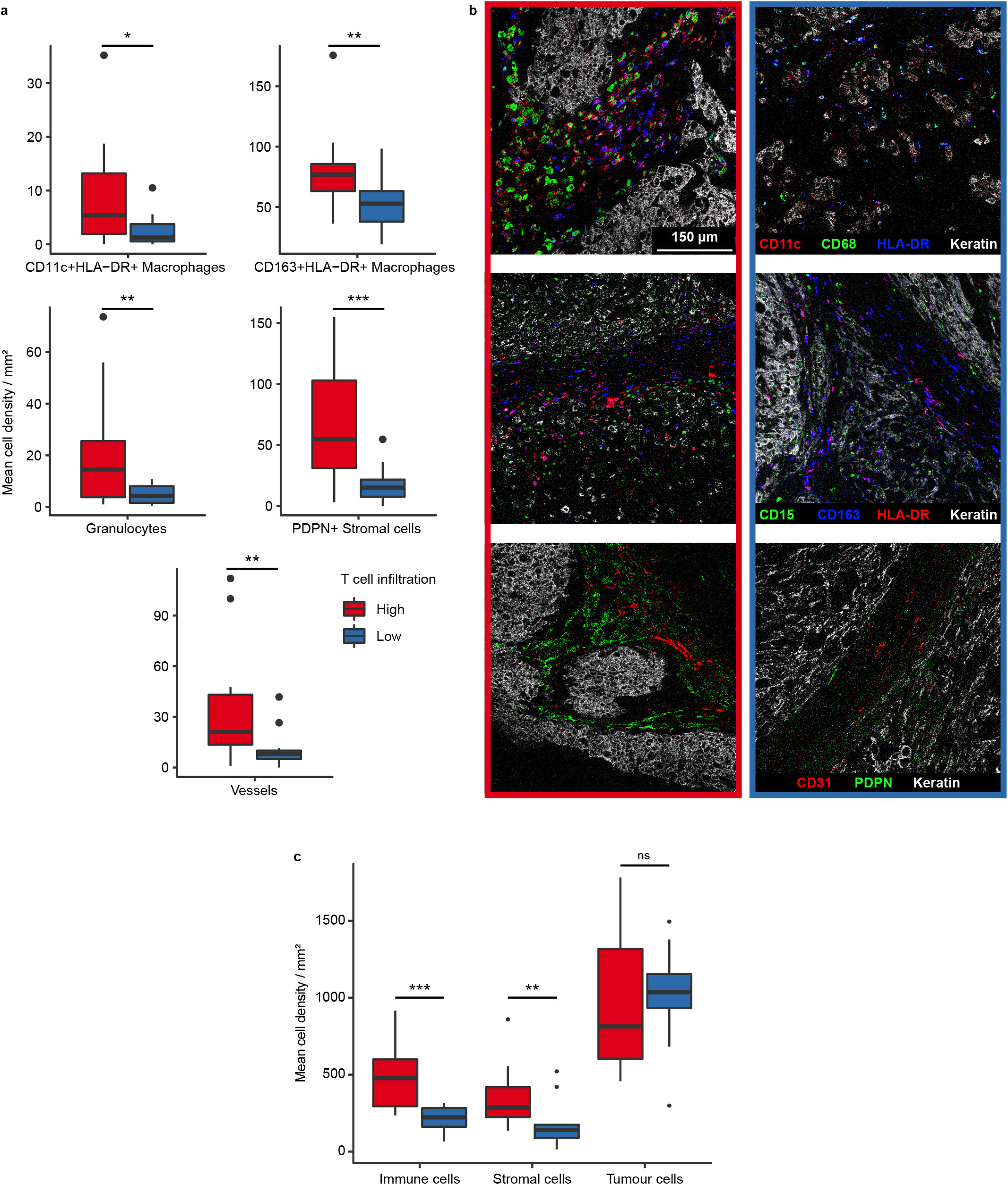
Comparative overview of the chordoma immune microenvironment. **a,** Boxplots with the significantly different phenotypes between the T cell high and T cell low groups. Two-sided t-test was performed in R and the *P* value was adjusted with the Benjamini-Hochberg false discovery rate (FDR). All presented phenotypes were significantly different between the T cell high an T cell low group. * = *P* < 0.05; ** = *P* < 0.01; *** *P* < 0.001. **b**, Representative images generated with imaging mass cytometry displaying the differentially abundant cell phenotypes in both T cell low and T cell high samples. Images are grouped per row for the displayed cell markers and outlined with their respective T cell group (T cell high = red rectangle & left, T cell low = blue rectangle & right). The top row presents CD11c^+^ macrophages (CD11c = red, CD68 = green, HLA-DR = blue, keratin = white), the middle row displays granulocytes and CD163^+^HLA-DR+ macrophages (CD15 = green, CD163 = blue, HLA-DR = red, keratin = white) and the bottom row shows vessels and podoplanin^+^ stromal cells (CD31 = red, PDPN = green, keratin = white). **c**, Boxplots presenting the cumulative mean cell density per mm^2^ of all immune cells, stromal cells and tumour cells in both T cell high and T cell low groups. Two-sided t-test was performed in R. ** = *P* < 0.01, *** = *P* < 0.001. PDPN = podoplanin.

**Supplementary Fig. 6.**
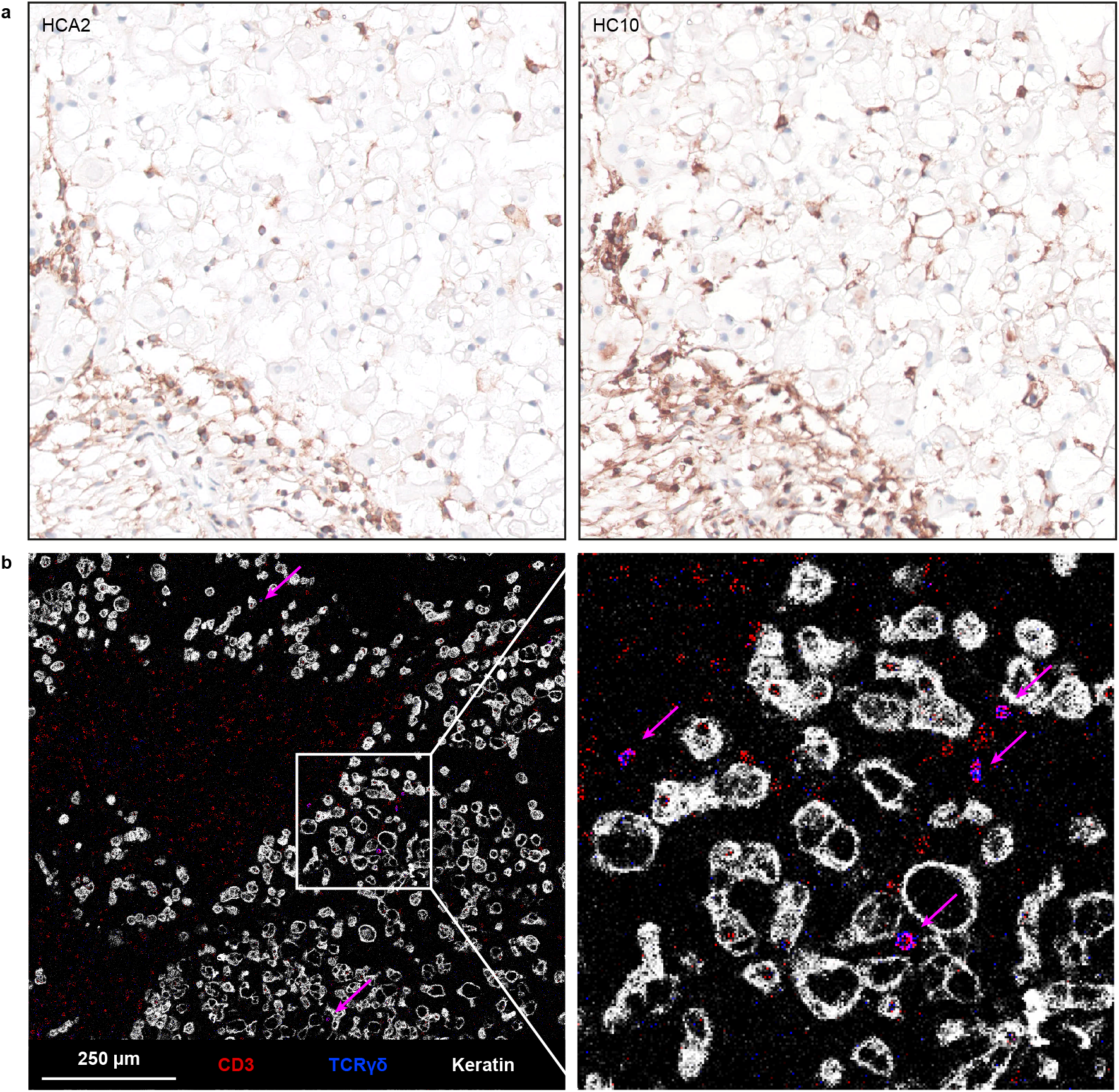
γδ T cell intiltration in relation to HLA class I expression. **a**, Examples images of Human Leukocyte Antigen (HLA) class I (HCA2 and HC10) expression in a T cell high chordoma as validated by immunohistochemistry. **b**, **A**n example image and a corresponding magnitication of the same tissue core as displayed in (**a**) as generated by imaging mass cytometry. The images present intratumoral γδ T cell intiltration as indicated by the magenta arrows (CD3 = red, TCRγδ = blue, keratin = white). TCR = T cell receptor.

**Supplementary Table 1. Top 200 most variably expressed genes across sarcoma subtypes**. Known chordoma-related genes are annotated in **bold**^1–3^. Genes are ordered following the heatmap presented in **Fig 1A**.

**Supplementary Table 2. Differentially expressed genes comparing chordomas with other sarcomas**. Contains the results from the differential gene expression analysis with DESeq2. Genes with a log2 fold change > 2 and an adjusted P < 0.01 were considered significantly differentially expressed. Known immune-related genes are annotated in the column “hgnc_symbol” with a yellow-filled cell. lfcSE = log fold change standard error; FDR = false discovery rate.

**Supplementary Table 3.** Univariate and multivariate cox proportional hazard results in the combined cohort. Table includes the results of the disease-specific, recurrence-free and metastasis-free survival. Significant parameters from the univariate analysis were taken along for the multivariable analysis. Patients who were only treated with radiotherapy were excluded from the recurrence-free survival analysis. Log-rank *P* values < 0.05 were considered significant and are annotated in bold and by significance level; * < 0.05, ** < 0.01, *** < 0.001. HR = hazard ratio; CI = confidence interval; RT = radiotherapy.

| Disease-specific survival |  |  | Metastasis-free survival |  |  |
| --- | --- | --- | --- | --- | --- |
| Variable | Univariate |  | Variable | Univariate |  |
|  | HR (95% CI) | log-rank *P* |  | HR (95% CI) | log-rank *P* |
| Age at diagnosis | 1 (0.98-0.99) | 0.2 | Age at diagnosis | 1 (0.99-1) | 0.3 |
| Max tumour size | 1 (0.91-1.2) | 0.7 | Max tumour size | 1.1 (0.95-1.2) | 0.3 |
| Location (Skull base)<sup>a</sup> |  |  | Location (Skull base)<sup>a</sup> |  |  |
| Mobile Spine | 0.56 (0.2-1.6) | 0.09 | Mobile Spine | 0.84 (0.21-3.4) | 0.8 |
| Sacrococcygeal | 0.35 (0.13-0.92) |  | Sacrococcygeal | 1.2 (0.4-4) |  |
| Treatment (RT alone) |  |  | Treatment (RT alone) |  |  |
| Surgery | 0.87 (0.24-3.1) | 0.6 | Surgery | 1 (0.23-5.1) | 1 |
| Surgery + RT | **0.57 (0.15-2.2) \*** |  | Surgery + RT | 0.95 (0.2-4.5) |  |
| Timing of RT (Adjuvant)<sup>b</sup> |  |  | Timing of RT (Adjuvant)<sup>b</sup> |  |  |
| Neoadjuvant | 0.66 (0.076-5.8) | 0.7 | Neoadjuvant | 3.6e-9 (0-Inf) | 0.2 |
| Surgical margin (R0)<sup>c</sup> |  |  | Surgical margin (R0) |  |  |
| R1 | 2.1 (0.23-19) | 0.3 | R1 | 2.6 (0.29-23) | 0.4 |
| R2 | 2.3 (0.75-6.8) |  | R2 | 0.68 (0.25-1.9) |  |
| T cell infiltration (High) |  |  | T cell infiltration (High) |  |  |
| Low | 2 (0.78-5.3) | 0.1 | Low | 1.5 (0.52-4) | 0.5 |
| Disease recurrence (No)<sup>c</sup> |  |  | Disease recurrence (No)<sup>c</sup> |  |  |
| Yes | 2 (0.78-5.3) | 0.9 | Yes | 6.3 (0.83-48) | **0.04 \*** |

| Recurrence-free survival<sup>c</sup> |  | Univariate |  | Multivariate |  |
| --- | --- | --- | --- | --- | --- |
| Variable |  | HR (95% CI) | log-rank *P* | HR (95% CI) | log-rank *P* |
| Age at diagnosis |  | 1 (0.98-1.02) | 0.8 |  |  |
| Max tumour size |  | 0.99 (0.9-1.1) | 0.9 |  |  |
| Location (Skull base)<sup>a</sup> |  |  |  |  |  |
| Mobile Spine |  | 1.95 (0.94-4.02) | **0.003 \*\*** | **2.2 (1-4.5) \*** | **0.01 \*** |
| Sacrococcygeal |  | 0.57 (0.26-1.2) |  | 0.81 (0.32-2.1) |  |
| Treatment (Surgery alone) |  |  |  |  |  |
| Surgery + RT |  | 0.82 (0.45-1.5) | 0.5 |  |  |
| Timing of RT (Adjuvant)<sup>b</sup> |  |  |  |  |  |
| Neoadjuvant |  | 0.29 (0.066-1.3) | 0.08 |  |  |
| Surgical margin (R0) |  |  |  |  |  |
| R1 |  | 1.6 (0.34-7.4) | 0.07 | 1.2 (0.24-5.6) |  |
| R2 |  | **2.3 (1.1-4.8) \*** |  | 1.7 (0.7-4.2) |  |
| T cell infiltration (High) |  |  |  |  |  |
| Low |  | 1.1 (0.6-2.1) | 0.7 |  |  |

<sup>a</sup>excluding the only patient with an extra-axial chordoma; <sup>b</sup>excluding patients who received a sandwich regimen of radiotherapy or radiotherapy alone; <sup>c</sup>excluding patients who were only treated with radiotherapy

**Supplementary Table 4.** Imaging mass cytometry marker panel. Ab = antibody, ON = overnight, PDPN = podoplanin.

|  | Target | Clone | Metal | Incubation Time | Temp | Dilution (x) |
| --- | --- | --- | --- | --- | --- | --- |
| Lymphoid | CD103 | EPR4166(2) | 168 Er | 5h | RT | 50 |
|  | CD20 | H1 | 142 Nd | Overnight | 4C | 100 |
|  | CD3 | EP449E | 153 Eu | Overnight | 4C | 50 |
|  | CD38 | EPR4106 | 169 Tm | Overnight | 4C | 100 |
|  | CD4 + 2nd AB | EPR6855 | 145 Nd | Indirect ON | 4C | 100 |
|  | CD7 | EPR4242 | 174 Yb | 5h | RT | 100 |
|  | CD8a | D8A8Y | 146 Nd | 5h | RT | 50 |
|  | FOXP3 | D608R | 159 Tb | Overnight | 4C | 50 |
| | TCR$\gamma\delta$ + 2nd Ab | H41 | 148 Nd | Indirect ON | 4C | 50 |
| Myeloid | CD11b | D6X1N | 144 Nd | 5h | RT | 100 |
|  | CD11c | EP1347Y | 176 Yb | 5h | RT | 100 |
|  | CD14 | D7A2T | 163 Dy | 5h | RT | 100 |
|  | CD15 | MC480 | 171 Yb | Overnight | 4C | 100 |
|  | CD163 | D6U1J | 173 Yb | 5h | RT | 50 |
|  | CD204 | J5HTR3 | 164 Dy | 5h | RT | 50 |
|  | CD68 | D4B9C | 143 Nd | Overnight | 4C | 100 |
|  | HLA-DR | TAL 1B5 | 141 Pr | 5h | RT | 100 |
| Stroma | CD31 | 89C2 | 147 Sm | Overnight | 4C | 100 |
|  | CD39 | EPR20627 | 157 Gd | 5h | RT | 100 |
|  | CD45 | D9M8I | 149 Sm | Overnight | 4C | 50 |
|  | CD45RO | UCHL1 | 165 Ho | Overnight | 4C | 100 |
|  | D2-40 (PDPN) | D2-40 | 166 Er | Overnight | 4C | 100 |
| | TGF-β | TB21 | 115 In | 5h | RT | 100 |
|  | Vimentin | D21H3 | 194 Pt | Overnight | 4C | 50 |
| Activation | Cleaved Caspase | 5A1E | 172 Yb | 5h | RT | 100 |
|  | Granzyme B | D6E9W | 150 Nd | 5h | RT | 100 |
|  | ICOS | D1K2T(TM) | 161 Dy | 5h | RT | 50 |
|  | IDO | D5J4E(TM) | 162 Dy | Overnight | 4C | 100 |
|  | Ki-67 | 8D5 | 152 Sm | Overnight | 4C | 100 |
|  | LAG-3 | D2G40(TM) | 155 Gd | 5h | RT | 50 |
|  | PD-1 | D4W2J | 160 Gd | 5h | RT | 50 |
|  | PD-L1 | E1L3N(R) | 156 Gd | Overnight | 4C | 50 |
|  | Tbet | 4B10 | 170 Er | 5h | RT | 50 |
|  | TIM-3 | D5D5R(TM) | 154 Sm | 5h | RT | 100 |
|  | VISTA | D1L2G(TM) | 158 Gd | 5h | RT | 100 |
| Tumour | CD56 | E7X9M | 167 Er | 5h | RT | 100 |
|  | CD57 | HNK-1 / Leu-7 | 151 Eu | Overnight | 4C | 100 |
|  | Keratin | C11 and AE1/AE3 | 198 Pt | Overnight | 4C | 50 |
|  | P16ink4a | D3W8G | 175 Lu | Overnight | 4C | 100 |
| | β-Catenin | D10A8 | 89 Y | Overnight | 4C | 100 |
| DNA | Histone H3 | D1H2 | 209 Bi | Overnight | 4C | 50 |

**Supplementary Table 5.**
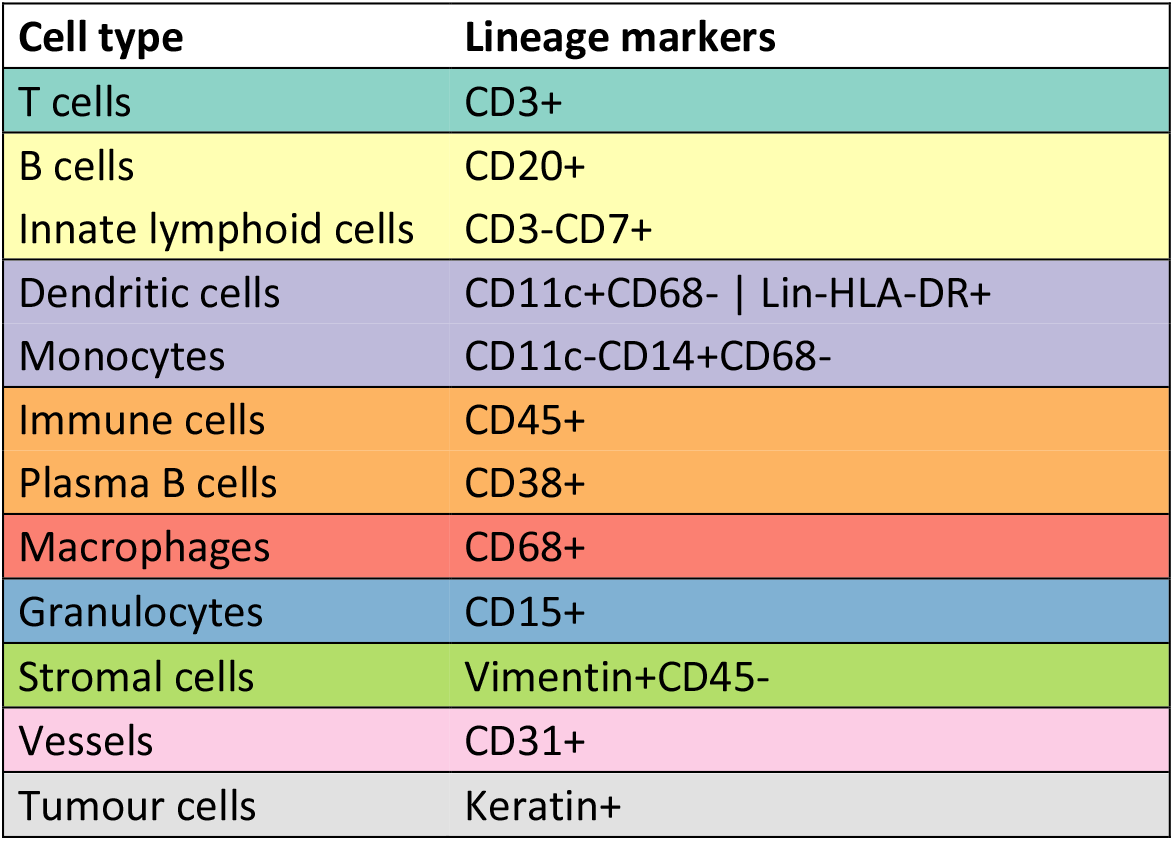
Cell type lineage markers imaging mass cytometry. Colours are indicating cell types according to overview heatmap in **Supplementary Fig 2**. Lineage markers are describing the used markers for cell type identification as used in this study. Lin = lineage markers e.g. CD14 and CD68.

